# Auditory localization improves with aligned saccadic orienting

**DOI:** 10.64898/2026.09.11.749382

**Authors:** Katarzyna Jurewicz, Krzysztof Basiński, Paweł Basoń, Marcin Leszczyński

**Affiliations:** Centre for Cognitive Science, Faculty of Philosophy, Jagiellonian University, Kraków, Poland; Auditory Neuroscience Laboratory, Department of Psychology, Medical University of Gdańsk, Gdańsk, Poland; Doctoral School in the Social Sciences, Jagiellonian University, Kraków, Poland; Department of Psychiatry, Columbia University College of Physicians and Surgeons, New York, USA

## Abstract

Animals actively sample the environment through movements of their sensory organs, yet whether these motor behaviors shape perception beyond the modality they directly control remains unresolved. In the auditory system, eye movements modulate neural activity from the auditory periphery to cortex, but their perceptual consequences have remained elusive. Here, we tested whether spontaneous eye movements exert a general influence on auditory perception or selectively interact with auditory computations involved in spatial orienting. Across four psychophysical experiments, participants freely explored natural scenes while performing auditory localization or pitch discrimination tasks. Brief sounds rapidly reorganized spontaneous gaze behavior, preferentially promoting large orienting saccades toward the sound source while suppressing saccades directed away. Critically, auditory localization systematically depended on the direction of spontaneous saccades: sound localization improved when saccades were directed toward the sound source and declined when saccades were directed away. No comparable relationship was observed during pitch discrimination despite closely matched auditory stimulation and visual exploration. These findings identify a selective coupling between saccadic behavior and auditory localization, providing a behavioral counterpart to the widespread oculomotor modulation observed throughout the auditory system. More broadly, our results suggest that the behavioral consequences of eye movements extend beyond vision, selectively engaging sensory computations linked to spatial orienting.

## Introduction

Animals actively control the sensory information they acquire. Rather than passively receiving inputs from the environment, they continuously determine what information is sampled and when it is processed by coordinating movements of their sensory organs with ongoing behavior. This principle, known as active sensing, is particularly evident in primates, where visual sampling is organized around rapid saccadic eye movements that redirect the fovea several times each second toward behaviorally relevant locations^1–4^. Beyond determining what information is sampled, accumulating evidence demonstrates that saccades also reorganize neural activity across widespread cortical and subcortical networks^5–14^. Within the visual system, this neural reorganization is accompanied by coordinated changes in visual processing, including saccadic suppression, predictive remapping of receptive fields, and presaccadic shifts of spatial attention^15–19^. Importantly, these changes to visual processing are feature-specific rather than global. Sensitivity to changes in motion and luminance is strongly altered across the saccade–fixation cycle, whereas other aspects of visual perception remain comparatively stable^16,20^. Thus, active vision is organized not by uniform sensory gain modulations, but by the selective prioritization of features and computations that are most relevant for ongoing behavior^14,21–24^.

Whether eye movements have a similar feature-selective influence on sensing beyond vision remains unclear. The widespread distribution of saccade-related signals raises the possibility that eye movements reorganize sensory computations in other modalities as well. Consistent with this idea, converging neurophysiological evidence demonstrates that eye movements modulate neural activity throughout the auditory hierarchy. Eye movement-related eardrum oscillations indicate that auditory processing is influenced already at the earliest stages of sound transduction^25–28^. Gaze position and saccades alter also neuronal activity in the auditory midbrain, thalamus, and auditory cortex in both humans and non-human primates^13,29–31^. Beyond these physiological effects, oculomotor behavior tracks multiple dimensions of auditory attention and expectation: eye movements reflect prioritized acoustic features during natural speech^32^, microsaccade direction tracks the spatial allocation of auditory attention^33^, and oculomotor dynamics reflect top-down attention and temporal expectations about upcoming sounds^34,35^. More broadly, auditory processing and motor behavior appear to be reciprocally linked. Motor activity can shape the temporal allocation of auditory attention, while auditory expectations can in turn organize ongoing oculomotor behavior^34,36,37^. Together, these findings demonstrate extensive interactions between auditory and oculomotor systems, although their computational and behavioral significance remains poorly understood^38^.

Yet the perceptual consequences of this widespread physiological modulation remain surprisingly unclear. Some aspects of auditory perception appear remarkably stable around eye movements: auditory detection and pitch discrimination are largely unaffected by saccades^39,40^. Other findings point to interactions with auditory spatial processing. Auditory judgments can improve when eye movements are directed toward the sound source^41^, auditory cues receive greater weight during audiovisual localization around saccades when visual information becomes less reliable^42^, and recent work has shown that sound lateralization depends on saccade direction^43^. These heterogeneous behavioral effects contrast with the widespread physiological influence of eye movements throughout the auditory system and suggest that the critical question may not be whether saccades influence auditory behavior, but which auditory computations are coupled to them.

One possibility is that eye movements exert a general influence on auditory perception that should be detectable across auditory features. Alternatively, their behavioral consequences may be selective, emerging primarily for auditory computations that are functionally linked to orienting behavior while leaving other aspects of hearing relatively unaffected. Spatial hearing provides a particularly informative test of this possibility. Unlike vision, which samples the environment most effectively in the direction of gaze, audition continuously monitors events beyond the current direction of gaze and provides spatial information that can be used to redirect attention and behavior toward relevant events. A fundamental function of auditory localization is therefore to determine where such events occur and support orienting toward them^44–48^. By contrast, non-spatial acoustic attributes such as pitch can be discriminated without determining where a sound originated. If the behavioral consequences of saccades are organized according to the functional role of auditory computations, their relationship with eye movements should therefore be particularly pronounced for localization rather than auditory perception in general.

This functional distinction is reflected in the neural organization of the auditory system. Spatial hearing recruits distributed networks linking auditory cortex with multisensory, attentional, and oculomotor circuits, whereas pitch processing relies predominantly on computations within the ascending auditory pathway and auditory cortex^49–55^. Auditory localization and pitch discrimination therefore provide a principled comparison for determining whether spontaneous eye movements interact selectively with auditory computations involved in spatial orienting rather than producing a general change in auditory performance. Critically, previous studies have typically examined different auditory features using different tasks, stimuli, and behavioral contexts, making this selectivity difficult to establish directly.

Here, we tested this hypothesis across four psychophysical experiments in which participants freely viewed natural scenes while performing near-threshold auditory judgments. Three experiments required localization of brief binaural or monaural sounds, whereas a fourth required discrimination of pitch using closely matched auditory stimulation and the same natural visual exploration. This design allowed us to directly compare the relationship between spontaneous saccades and spatial versus non-spatial auditory performance under closely matched behavioral conditions. Brief sounds rapidly reorganized ongoing eye movements, preferentially promoting large saccades toward the sound source. Critically, saccadic behavior selectively facilitated auditory localization: performance improved when saccades were directed toward the sound and declined when they were directed away, whereas no comparable directional relationship emerged during pitch discrimination. Together, these findings reveal a selective coupling between saccadic responses and auditory localization and suggest that the behavioral consequences of eye movements extend beyond vision by preferentially engaging sensory computations linked to spatial orienting.

## Results

Participants freely viewed natural scenes while performing one of four near-threshold auditory discrimination tasks (Figure 1). During each 8-s viewing period, one or two brief auditory probes were presented while participants explored the scene with unrestricted eye movements. Three experiments required spatial judgments about the sound source, using either binaural interaural time differences (Experiments 1a and 1b) or monaural stimulation (Experiment 2), whereas Experiment 3 required discrimination of a non-spatial acoustic feature (pitch). This design allowed us to compare the relationship between saccadic eye movements and auditory perception across tasks that either did or did not require spatial localization.

**Figure 1.**
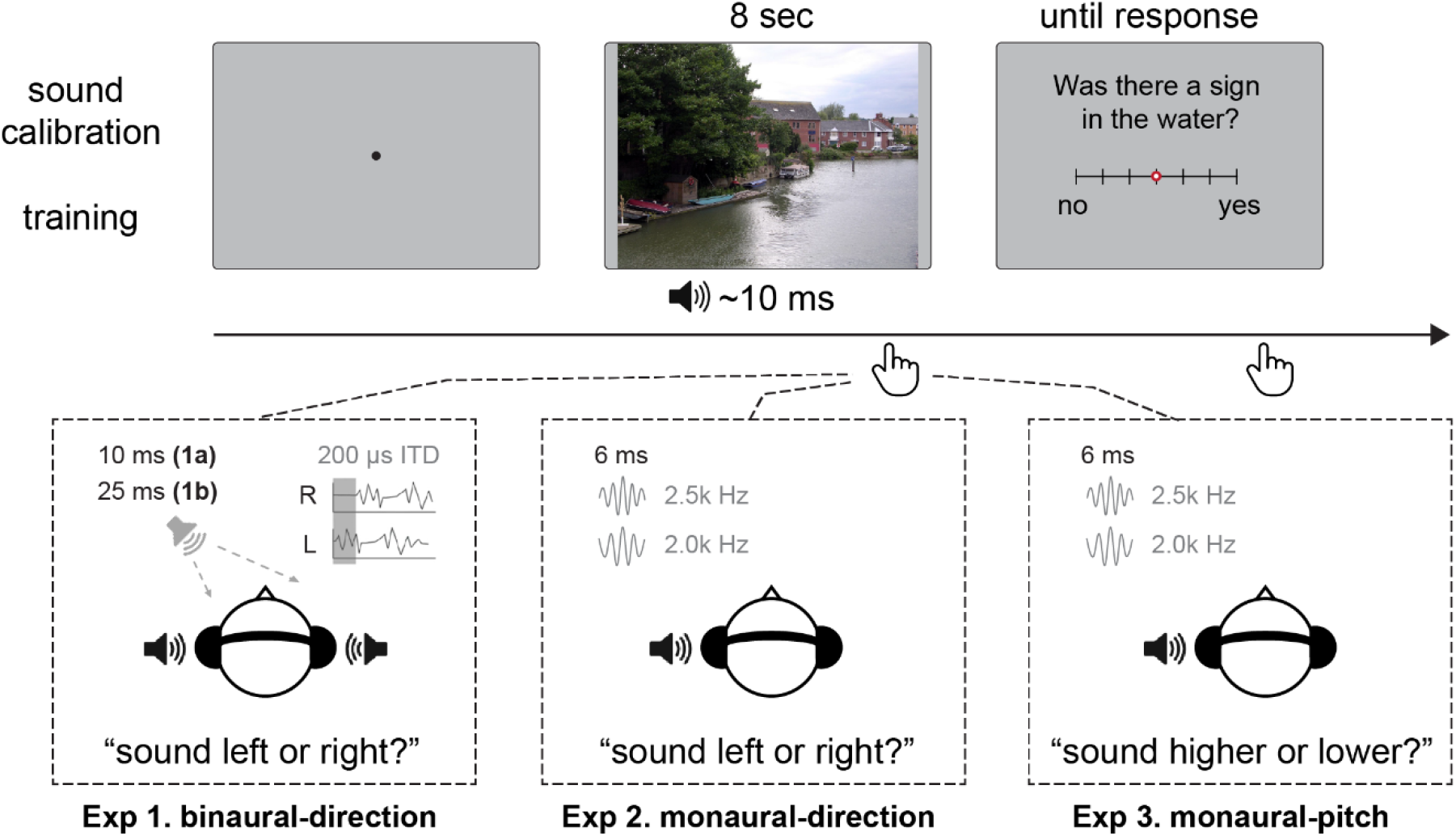
Study design and experiments. Participants performed an auditory discrimination task while freely viewing natural scenes. As illustrated in the trial timeline (top), each trial began with a fixation screen, followed by an 8-second scene presentation during which brief auditory probes (one or two) were delivered via headphones. After scene offset, participants rated their agreement with a scene content probe (e.g., “Was there a sign in the water?”) on a scale from “no” to “yes,” with no time limit. The experiments (bottom) differed in auditory stimulus type and task: Experiment 1 (binaural-direction) presented brief pink noise (10 ms in Experiment 1a and 25 ms in Experiment 1b) binaurally with a small interaural time delay (ITD), and participants judged whether the sound originated from the left or right; Experiment 2 (monaural-direction) presented a brief (6 ms) pure tone to one ear (either 2 or 2.5 kHz), with participants performing the same left/right laterality judgment; Experiment 3 (monaural-pitch) used the same monaural tone as Experiment 2 but required pitch discrimination (higher or lower). The auditory stimulus volume was individually calibrated prior to the main experiments, and the experiments were preceded by the training.

### Auditory stimuli rapidly reorganize spontaneous saccades

We first asked whether brief auditory stimuli influence spontaneous eye movements during natural visual exploration. To this end, we quantified the temporal distribution of saccades relative to sound onset (Figure 2). Saccade counts were expressed as changes relative to the 500 ms interval preceding sound presentation, which served as the baseline. Brief auditory stimuli elicited a rapid directional reorganization of spontaneous saccades. Across all four experiments, saccades directed away from the sound were suppressed immediately following sound onset for about 600 ms, the time before the task response (peak suppression, Experiment 1a: 375 ms; Experiment 1b: 300 ms; Experiment 2: 250 ms; Experiment 3: 300 ms), whereas saccades directed toward the sound recovered substantially faster after the initial suppression (Figure 2; lack of significant difference from baseline, Experiment 1a: 275 ms; Experiment 2: 225 ms; Experiment 3: 225 ms). The suppression of sound-incongruent saccades was remarkably consistent across experiments. Relative to baseline, the number of incongruent saccades reached a minimum of 25% below baseline in Experiment 1a (Figure 2A), 31% below baseline in Experiment 1b (Figure 2B), 36% below baseline in Experiment 2 (Figure 2C), and 27% below baseline in Experiment 3 (Figure 2D; all *p* < 0.001, Wilcoxon signed-rank tests, Benjamini–Hochberg corrected). Consequently, the number of sound-congruent saccades exceeded the number of sound-incongruent saccades over extended post-stimulus intervals in all four experiments (Experiment 1a: 125–450 ms; Experiment 1b: 250–375 ms; Experiment 2: 150–500 ms; Experiment 3: 100–575 ms; all *p* < 0.001, Wilcoxon tests, Benjamini–Hochberg corrected).

**Figure 2.**
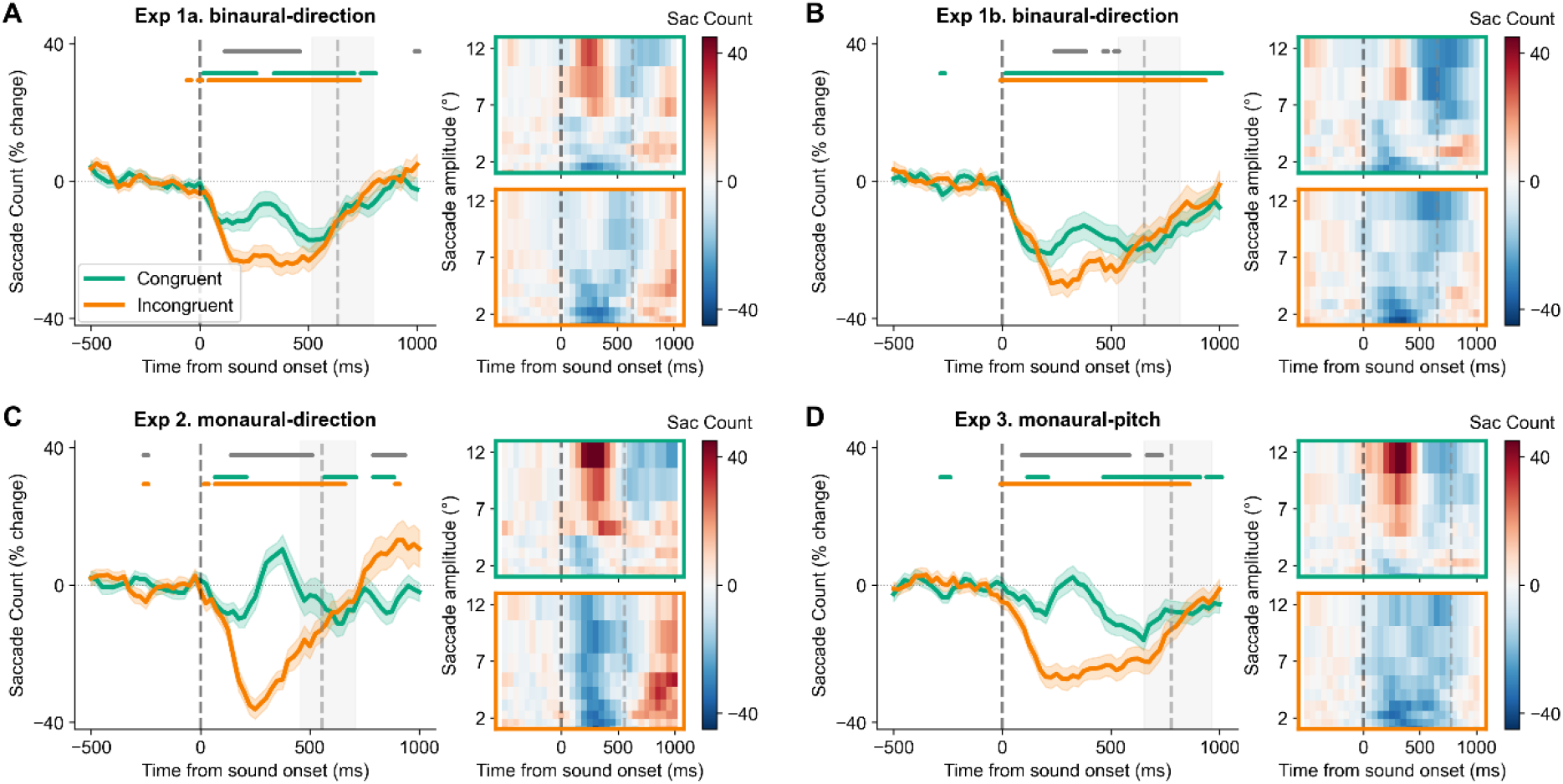
Saccadic behavior after sound. A) Left: Saccade count in Experiment 1a displayed as a percent change from pre-sound baseline, for saccade onsets happening at different intervals from the sound onset, separately for saccades with congruent (green line) or incongruent (orange line) direction to the sound direction. Shaded areas represent SEM across participants. Horizontal bars show significant, FDR-corrected difference from baseline for congruent (orange) and incongruent (green) saccades, and the difference between the relative count of congruent and incongruent saccades (gray). Black vertical dashed line indicates sound onset, gray vertical dashed line marks average manual response time in the auditory task, gray shading marks the 25%–75% range of response times. Right: Saccade count (percent change from pre-sound baseline) at different intervals from the sound onset, split for different saccade amplitudes, separately for congruent (orange frame) and incongruent (green frame) saccades. Red color indicates increases, blue color indicates decreases. Ten amplitude bins were obtained by quantile binning amplitudes of the saccades in the baseline period. The same conventions were used for displaying the saccade count in B) Experiment 1b, C) Experiment 2, D) Experiment 3.

We next asked whether this saccadic response differs in saccade amplitude. Large-amplitude saccades (> approximately 5° visual angle) showed a pronounced directional bias toward the sound, whereas small-amplitude saccades were uniformly suppressed regardless of direction (Figure 2, heatmaps). In all experiments, the number of large sound-congruent saccades increased following sound presentation, whereas large saccades directed away from the sound exhibited only a weak increase (Experiments 1a and b) or remained suppressed (Experiments 2 and 3). This interaction between saccade direction and amplitude reached significance in Experiments 2 and 3 following correction for multiple comparisons (175–350 ms and 200–375 ms, respectively; repeated-measures ANOVA, all *p* < 0.05, Benjamini–Hochberg corrected; Supplementary Figure S1) and showed the same qualitative pattern in Experiments 1a and 1b. Together, these findings demonstrate that brief auditory stimuli do not simply suppress spontaneous eye movements. Instead, they rapidly reorganize ongoing oculomotor behavior by selectively promoting large saccades toward the sound while suppressing saccades directed away. Whether this rapid auditory-driven reorganization of spontaneous saccades modulates auditory perception remained unknown. We therefore next asked whether auditory performance depended on the direction of the accompanying saccade.

### Auditory localization is selectively facilitated by spontaneous saccades

We next asked whether the auditory-driven reorganization of spontaneous saccades was accompanied by changes in auditory performance. To address this question, we quantified auditory localization and pitch discrimination accuracy as a function of spontaneous saccade direction relative to the sound source (Figure 3). Auditory performance associated with saccades occurring during the 500 ms preceding sound onset served as the baseline. In all three localization experiments (Exp. 1a, 1b and 2), auditory performance depended on the direction of the accompanying saccade. Responses were significantly more accurate when participants generated spontaneous saccades toward the sound than when they generated saccades directed away from it. Following sound-congruent saccades, localization accuracy increased above baseline in both binaural (Experiments 1a and b) and monaural (Experiment 2) localization tasks, whereas accuracy following sound-incongruent saccades decreased below baseline (Experiment 1a: congruent, 275–425 ms; incongruent, 200–450 ms; Experiment 1b: congruent, 475 ms; incongruent, 200–525 ms; Experiment 2: congruent, 275–300 ms; incongruent, 125–425 ms; all *p* ≤ 0.04, Wilcoxon signed-rank tests, Benjamini– Hochberg corrected). Consequently, localization performance was reliably higher following congruent than incongruent saccades in all three experiments (Experiment 1a: 200–500 ms; Experiment 1b: 225–500 ms; Experiment 2: 200–425 ms; all *p* < 0.001; Figure 3A-C).

**Figure 3.**
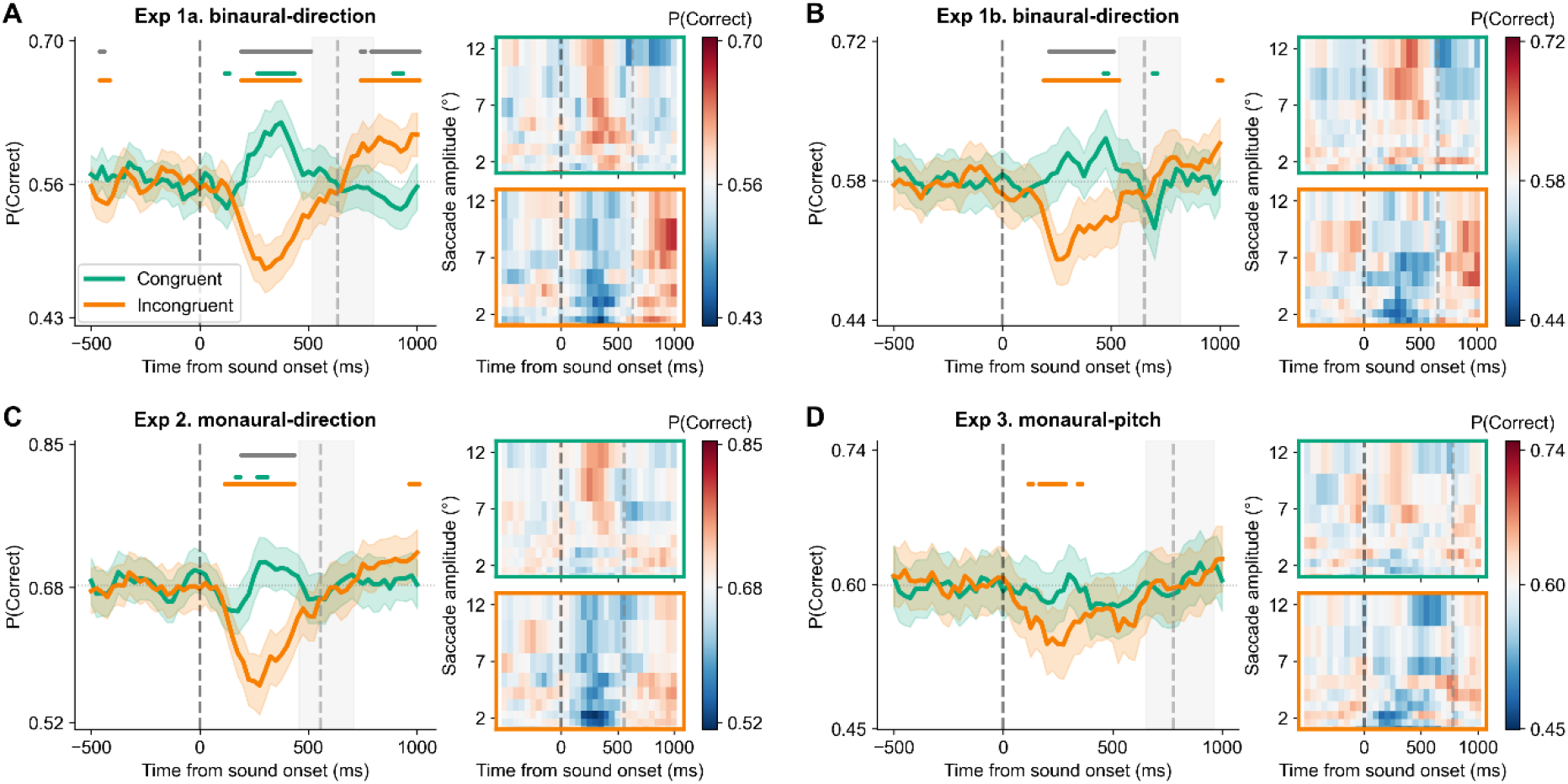
Performance in auditory tasks aligns with saccadic responses. A) Left: Probability of a correct response in the Experiment 1a. auditory task, for saccade onsets happening at different intervals from the sound onset, separately for saccades with congruent (orange line) or incongruent (green line) direction to the sound side. The Y axis was calibrated to show the mean and +/-25% of the mean probability value range, based on correctness conditioned on the pre-sound saccades. Shaded areas represent SEM across participants. Horizontal bars show significant, FDR-corrected difference from baseline for congruent (green) and incongruent (orange) saccades, and the difference between correctness following congruent and incongruent saccades (gray). Black vertical dashed line indicates sound onset, gray vertical dashed line marks average manual response time in the auditory task, gray shading marks the 25%–75% range of response times. Right: Probability of a correct response for saccades at different intervals from the sound onset, split for different saccade amplitudes, separately for congruent (orange frame) and incongruent (green frame) saccades. Red color indicates increases, blue indicates decreases. Ten amplitude bins were obtained by quantile binning amplitudes of the saccades in the baseline period. The same conventions were used for displaying probability of a correct response in B) Experiment 1b, C) Experiment 2, D) Experiment 3.

Importantly, this relationship disappeared when the task no longer required spatial processing. In the pitch discrimination experiment (Experiment 3) only sound-incongruent but not sound-congruent saccades significantly altered perceptual accuracy relative to baseline (Experiment 3: incongruent, 125–275 ms; p < 0.001), and no reliable difference between saccade directions was observed (Figure 3D). Despite identical visual exploration and highly similar auditory stimulation, saccades selectively benefited auditory localization tasks while leaving discrimination of a non-spatial acoustic feature unaffected. Reaction times mirrored the accuracy results. Participants responded significantly faster following sound-congruent than sound-incongruent saccades in all three localization experiments, whereas no such advantage was observed during pitch discrimination (Supplementary Figure S2). Expressing localization performance as signal-detection sensitivity (*d′*) reproduced the same qualitative congruency pattern (Supplementary Figure S2), indicating that the observed effects were not attributable to differences in response frequency or response omissions. Together, these findings demonstrate that spontaneous saccades do not interact uniformly with auditory behavior. Instead, congruent saccades selectively facilitate auditory localization while leaving non-spatial auditory discrimination largely unaffected.

### Large saccades towards the sound facilitate auditory processing

Having established that auditory localization selectively benefited from spontaneous saccades directed toward the sound source, we next asked whether this behavioral advantage depended on the properties of the accompanying eye movement. Because the saccadic responses described above were dominated by large-amplitude saccades, we hypothesized that perceptual facilitation would likewise be strongest for large-amplitude movements. To test this prediction, we quantified auditory performance separately for different saccade amplitudes (Figure 3, heatmaps; see Supplementary Figure S3 for statistics).

Across all three localization experiments, improvements in auditory performance (above-baseline accuracy) were concentrated among large-amplitude saccades directed toward the sound source. In Experiment 1a, localization accuracy following sound-congruent saccades increased for saccade amplitude above 4° peaking around 5-7° (Figure 3A). A comparable pattern was observed in Experiments 1b and 2, peaking around 7-10° (Figure 3B,C). Impairment in auditory performance (below-baseline accuracy) was in turn concentrated among small-amplitude saccades incongruent with the sound location. Performance following sound-incongruent saccades remained lower across amplitude bins, exhibiting lowest values for saccades at or below 2°. The effect of amplitude reached significance in Experiments 1b and 2 following correction for multiple comparisons (250, 300–325 ms and 275–375, 425–450 ms, respectively; repeated-measures ANOVA, all *p* < 0.05, Benjamini– Hochberg corrected; Supplementary Figure S3) and showed the same qualitative pattern in Experiment 1a. Consequently, larger saccades supported better localization performance in addition to the congruency effect. In the pitch discrimination experiment (Experiment 3), larger saccades were also coupled with better performance, i.e. larger saccades tended to support better performance despite the lack of congruency effect, although the amplitude effect was not significant after correction for multiple comparisons (Supplementary Figure S3).

These findings indicate that auditory performance is coupled to spontaneous saccadic responses, with the strongest behavioral advantage accompanying large saccades. Across all experiments, auditory stimuli rapidly reorganized saccadic responses, and this reorganization was associated with improved auditory perception during natural visual exploration. This behavioral advantage was strongest for large saccades directed toward the sound source in localization tasks, suggesting that the relationship between spontaneous eye movements and auditory perception is selective for spatial computations rather than reflecting a general modulation of auditory processing.

## Discussion

Eye movements modulate neural activity throughout the auditory system, yet behavioral work has produced surprisingly little evidence that they systematically influence auditory performance. Here, we show that the behavioral consequences of spontaneous saccades are selective. Brief sounds rapidly reorganized ongoing eye movements, increasing the probability of large saccades toward the sound source while suppressing saccades in the opposite direction. Critically, auditory localization improved when spontaneous saccades were directed toward the sound and declined when they were directed away, whereas no comparable directional relationship emerged during pitch discrimination despite closely matched sensory stimulation and visual behavior. Together, these findings provide a behavioral counterpart to the widespread oculomotor modulation observed throughout the auditory system and suggest that eye movements interact selectively with auditory computations involved in spatial localization rather than exerting a general influence on auditory perception.

### Eye movements selectively couple to spatial processing

Our findings show that spontaneous saccades are selectively coupled to auditory localization rather than to auditory perception in general. Localization performance systematically depended on whether saccades were directed toward or away from the sound source, whereas no comparable directional relationship emerged during pitch discrimination despite closely matched auditory stimulation and visual exploration. This selectivity suggests that the consequences of eye movements for sensory processing depend on the computations being performed rather than reflecting a uniform modulation of sensory sensitivity. A similar principle is well established in active vision, where saccades selectively reorganize visual processing according to the demands of ongoing behavior. Visual sensitivity is transiently reduced around saccades to limit the consequences of self-generated retinal motion^16,18,56^, whereas predictive remapping, presaccadic attentional allocation, and trans-saccadic integration support the selection and integration of information across successive fixations^15,17,19^. Together, these observations suggest that coupling between oculomotor behavior and sensory processing may represent a broader principle extending beyond the sensory modality directly controlled by the movement. Consistent with this possibility, oculomotor behavior has been linked to temporal expectation and perceptual performance even for tactile events^57^. This interpretation converges with evidence for reciprocal links between auditory and motor systems: motor activity can organize auditory processing in time, while auditory temporal expectations can organize oculomotor behavior^34,36,58^. Our findings reveal a complementary spatial relationship, in which auditory localization is selectively associated with ongoing saccadic orienting. Auditory localization provides spatial information that can guide attention, gaze, and action toward behaviorally relevant events^48^, whereas discriminating non-spatial features such as pitch does not necessarily require spatial orienting. Oculomotor signals may therefore preferentially interact with auditory computations that contribute to orienting rather than uniformly modulating auditory processing. From this perspective, the relevant distinction may not be between visual and auditory processing, but between sensory computations that are functionally linked to orienting and those that are not. Auditory localization may be preferentially coupled to eye movements precisely because both contribute to selecting where behavior should be directed next.

Natural orienting, however, involves coordinated movements of the eyes, head, and body and depends on the spatial and contextual relationship between auditory and visual information. The present experiments isolated eye movements during free visual exploration but therefore they capture only one component of this broader sensorimotor system. Future studies using more immersive environments and unrestricted orienting behavior will be important for determining how eye, head, and body movements jointly interact with auditory localization under natural conditions.

### Why auditory localization?

Auditory localization is particularly well positioned to interact with saccadic responses because hearing and vision make complementary contributions to spatial behavior. Unlike vision, which samples the environment most effectively in the direction of gaze, audition continuously monitors events both within and beyond the current field of view. Spatial hearing can therefore identify behaviorally relevant events outside the current direction of gaze and provide information that guides subsequent shifts of attention, gaze, and action; conversely, active orienting movements can themselves improve auditory localization^44– 48,54,59^. Coupling auditory spatial information to the systems controlling orienting could thus provide an efficient mechanism for coordinating sensory sampling across modalities.

This functional relationship is reflected in the organization of the auditory system. Spatial hearing recruits distributed circuits linking auditory cortex and auditory midbrain with multisensory, attentional, and oculomotor networks, including collicular pathways through which auditory information can directly guide spatial attention and orienting^50,52,60^, whereas non-spatial acoustic attributes such as pitch depend more strongly on computations within auditory pathways and auditory cortex^51,53^. Consistent with this organization, we observed a robust relationship between spontaneous saccades and localization across both binaural and monaural spatial tasks, but not during pitch discrimination. More broadly our findings suggest that the critical determinant of saccade-related modulation is not sensory modality itself, but the functional relationship between a sensory computation and orienting behavior. Importantly, pitch provides only one non-spatial comparison. Auditory scene analysis, speech perception, hearing in noise, intensity judgments, and other auditory computations may interact with eye movements in ways not captured by the present experiments. Identifying which processes are coupled to saccadic behavior will be important for establishing the generality of the proposed principle.

### From auditory physiology to auditory perception

Oculomotor signals are represented throughout the auditory system. Eye movement-related signals have been observed from the auditory periphery to the inferior colliculus, medial geniculate body, auditory cortex, and higher-order association areas^25,30,61,62^. Spontaneous saccades also reorganize cortical auditory processing by resetting ongoing oscillatory activity, modulating neuronal excitability, and altering functional interactions between auditory and oculomotor regions^13,31^. Yet the behavioral significance of these signals has remained difficult to establish. Auditory detection and pitch discrimination, for example, show little or no peri-saccadic modulation^39,40^.

The present results suggest one way to reconcile widespread physiological modulation with apparently limited behavioral effects. Oculomotor signals need not produce a uniform change in auditory sensitivity to be behaviorally relevant. Instead, their consequences may emerge selectively when auditory information contributes to computations shared with the orienting system. Under this account, the absence of measurable effects on detection or pitch discrimination is not inconsistent with extensive oculomotor modulation throughout the auditory hierarchy. Rather, neural modulation may become behaviorally expressed only for particular computations or behavioral contexts. Importantly, our experiments do not establish that the physiological effects reported previously directly generate the behavioral effects observed here. Nevertheless, the convergence is notable. The widespread availability of information about eye position and eye movements throughout the auditory system provides a potential substrate through which auditory representations could be coordinated with ongoing orienting behavior. Determining how these neural signals relate to the behavioral coupling identified here will require simultaneous measurements of auditory neural activity, eye movements, and perceptual decisions.

### What links spontaneous saccades and auditory localization?

The present results establish a relationship between saccadic behavior and auditory localization but do not determine its causal direction or processing stage. Several mechanisms could produce this relationship. One possibility is that auditory localization and saccadic behavior draw on a shared spatial decision variable. Auditory evidence about sound location and evolving oculomotor plans could be represented within partially overlapping spatial codes, such that orienting toward one side shifts the resulting auditory decision in the same direction. When auditory and oculomotor signals are aligned, their contributions would reinforce one another, producing the increased localization performance observed for sound-congruent saccades; when they are opposed, they would compete, reducing performance. Such an account explains why congruent and incongruent saccades have opposite consequences for localization accuracy. A second possibility is that auditory localization and saccadic response are parallel consequences of a shared sensorimotor computation. Rather than the eye movement influencing the auditory decision, auditory spatial evidence could feed into a common orienting representation that contributes both to gaze direction and to the eventual localization response. Stronger or more reliable spatial evidence would then increase both the probability of orienting toward the sound and the probability of reporting its location correctly. This account predicts covariation between gaze and auditory choice without requiring a causal effect of the eye movement itself. A third possibility is that oculomotor signals enter the auditory decision process after the initial sensory representation has been established. Under this account, saccades may retroactively refine or stabilize auditory spatial representations after the initial sensory percept has been formed. Alternatively, information about the saccade direction could bias the accumulation, maintenance, or readout of auditory spatial evidence during decision formation. These mechanisms are not mutually exclusive. Auditory spatial representations, orienting plans, decision variables, and response selection may interact continuously as behavior unfolds. The present findings therefore locate the effect within a broader perception–action transformation rather than assigning it exclusively to sensory encoding or motor output.

### Eye movements as global coordinating signals

The present findings broaden the functional significance of eye movements beyond visual sampling. A saccade changes what the visual system samples, but the associated motor signals are broadcast far beyond visual cortex. Our results suggest that one consequence of this widespread signaling is the selective coupling of eye movements to computations in other sensory systems that contribute to spatial behavior. In audition, this coupling is expressed most clearly during localization: performance is facilitated when auditory and oculomotor directions align and impaired when they conflict, while non-spatial pitch judgments show no comparable directional relationship. This perspective offers a framework for understanding why oculomotor signals are found throughout the auditory hierarchy despite their apparently modest effects on hearing in general. Their function may not be to globally enhance or suppress auditory processing. Instead, they may help align auditory information with the sensorimotor systems that determine where attention and behavior are directed next. More broadly, eye movements may constitute one component of a distributed control architecture through which sensory processing is coordinated with ongoing action.

## Methods

### Subjects

We aimed for a sample size of ≥40 participants per experiment. This number was chosen to exceed sample sizes used in all previous studies testing auditory performance around the saccade that were similar to ours (Harris and Lieberman^39^: 5 and 7 subjects in Exp 1-2; Rorden and Driver^41^: 10, 10, 8, and 14 subjects in Exp 1-4; Bröhl & Kayser^40^: 14 subjects; Sotero Silva et al.^43^: 34 subjects). In Experiment 1a (binaural-direction, 10 ms stimulus) data from N = 52 participants are reported (mean age = 22.8, SD = 2.4, 38 females). Two additional participants did not complete the session due to technical issues. In Experiment 1b (binaural-direction, 25 ms stimulus) data from N = 40 participants are reported (mean age = 22.7, SD = 4.01, 30 females). In Experiment 2 (monaural-direction) data from N = 40 participants are reported (mean age = 23.8, SD = 3.9, 32 females). In Experiment 3 (monaural-pitch) data from N = 46 participants are reported (mean age = 22.1, SD = 3.7, 36 females). All participants were naive to the aims of the experiment, reported normal hearing and no history of neurological disorders. The study was approved by the institutional review board at the Jagiellonian University, and all subjects gave written informed consent.

#### Experimental Setup

Recordings took place in a soundproofed booth with dim, controlled lighting. Participants remained seated in front of a table with their head on a chinrest, aligned to the center of a monitor 90 cm away (AOC 24P2W1DG5, 23.8’’, 1920 × 1080 px, 60 Hz refresh rate), which presented visual stimuli. Sounds were presented via headphones (Beyerdynamic DT 770 PRO 80 OHM). The experimental tasks were implemented in PsychoPy^63^.

Eye movements were recorded at 1000 Hz from the right eye using a desktop mounted infrared-based EyeLink 1000 Plus Eye Tracker (SR Research Ltd.) running Host Software version 5.50. Eye-tracking calibration and validation was performed for each block using the built-in 9-point grid.

#### Procedure

We conducted a series of psychophysical experiments, in which participants performed auditory tasks while freely viewing natural scenes. Each experiment followed a similar procedure: participants were first instructed about the auditory task and completed a threshold estimation procedure (see below). Then they were familiarized with the visual task and performed training trials on this task alone (3 scenes). Next, they performed training trials of the actual experiment, containing both auditory and visual stimuli (min 5 scenes, extended if needed), followed by ten blocks of the main experiment (20 scenes each).

#### Visual stimuli and task

Scenes in the experiments were 1920 × 1080 (spanning 32 × 18 degrees of visual angle [dva]) color photographs of indoor and outdoor environments, often featuring people and animals, sourced from the Nencki Affective Picture System (NAPS)^64^. Images were pre-selected according to two criteria: subjective high complexity (rated by the researcher) and neutral to slightly positive valence values.

During the experiment, each scene was presented for 8 s. Following each presentation, participants completed a scene content probe (e.g., “A boy had a yellow T-shirt”) to encourage active scene exploration; agreement was indicated on a 7-point scale from 1 (*disagree*) to 7 (*agree*). There was no time limit for this response.

#### Auditory stimuli and tasks

The auditory tasks were designed around each participant’s individual perceptual threshold. During each scene presentation, 1 or 2 auditory probes were delivered at randomly selected times via headphones (380-400 probes per experiment). Specifically, we always used 2 auditory probes per scene in Experiment 1a, and added 5% of “catch” sound probes (no sound) in Experiments 1b and 2, 3. The first sound could appear 1000 to 3500 ms after image onset, the second sound could appear 2500 ms after the first sound at the earliest and 1500 ms before image offset at the latest. All sounds were rendered offline as WAV files with a 48 kHz sample rate using REAPER v.7.11.

In Experiment 1a identical 10-ms pink noise was presented binaurally, while the interaural time difference (200 μs delay) was introduced to evoke perception of left-ward or right-ward sound origin. Participants reported sound direction via button press (index and middle finger of the right hand for “left” and “right” response respectively).

Experiment 1b replicated Experiment 1a with two minimal modifications: (1) the auditory stimulus was extended to 25 ms, to extend the results of Experiment 1 with a different auditory probe; (2) “catch” trials were included, i.e. we included scene presentations with a single auditory stimulus (instead of two auditory stimuli).

In Experiment 2. a 6-ms tone (a sinusoid of 2 or 2.5 kHz) was presented monaurally to either left or right ear. Participants reported sound direction via button press (index and middle finger of the right hand for “left” and “right” response respectively).

In Experiment 3. a 6-ms tone (a sinusoid of 2 or 2.5 kHz) was again presented monaurally to either left or right ear. This time, participants discriminated the sound’s pitch via button press (index and middle finger of the right hand for “lower” and “higher” response respectively).

#### Staircase auditory calibration

The 3-down 1-up staircase procedure was used to adjust the volume of auditory probes for each participant. After 3 consecutive correct responses, volume was decreased by a randomized amount that depended on the current volume level (larger drop for higher values). A single error increased volume by 10%. The procedure ended when accuracy, computed as a rolling average over the last 20 trials of each type, fell within the desired range (no earlier than trial 40). The volume at termination became the stimulus volume for the main task. In Experiment 1a, the accuracy for left-ward and right-ward stimulus had to each fall within [0.70, 0.75], and the starting volume was set as 0.0001 of the max. In Experiment 1b the accuracy for left-ward and right-ward stimulus had to each fall within [0.70, 0.75], and the starting volume was set as 0.0005 of the max. In Experiments 2 and 3 accuracy for high pitch and for low pitch (each computed across both ears) had to both fall within [0.70, 0.80], and accuracy for each of the four individual stimulus types (left-high, left-low, right-high, right-low) had to fall within [0.50, 1.00] each. Starting volume was 0.0005 of the max.

#### Data processing and statistical analyses

We used saccade extraction implemented by the EyeLink system (“cognitive” setting, a velocity threshold of 30°/s and an acceleration threshold of 8000°/s^2^). Eye positions were recorded as horizontal and vertical coordinates in degrees of visual angle.

We selected all saccades that were within -500 to 1000 ms from the sound onset (saccade could fall within this temporal distance to only one sound in the trial). We assigned the response associated with the sound to each saccade within this period, and defined congruency of saccade direction with sound direction based on starting and ending position of the saccade along the horizontal axis.

We performed the main analyses in a running window of 100 ms width and 25 ms step. We focused on two dependent variables, analyzed separately: (1) normalized saccade count (percentage change relative to a per-condition baseline of −500 to 0 ms), and (2) the proportion of correct responses. To measure the number of saccades at different intervals relative to sound onset, we summed the number of saccades in a given time window for all sounds, separately for both congruency conditions. Then, to normalize the values to an easily interpretable scale, we divided the number of saccades in each time window by the number of saccades in the baseline period (−500 to 0 ms from sound onset, adjusted for the difference of window length, i.e. divided by 5) within the condition. To measure auditory task performance depending on the moment of saccade execution, we calculated the fraction of correct responses associated with saccades in a given subject-window-congruency condition cell.

The statistical comparisons were two-fold. (1) We compared the number of saccades, and the probability of correct response with the values obtained in the baseline period separately for congruent and incongruent saccades. Similarly, we directly compared the values between these two congruency conditions. For these comparisons we used Wilcoxon signed-rank tests, false discovery rate (FDR) corrected for multiple comparisons using the Benjamini–Hochberg procedure (183 tests per correction: 3 pairwise comparisons × 61 time windows). (2) To include amplitude factor into the analyses of timing and congruency, we performed repeated measures ANOVAs. Saccade amplitude was split into three quantile-based bins per subject, where bin boundaries were determined exclusively on the pre-sound saccades (onset before sound presentation). This resulted in a 2 × 3 fully within-subjects design: congruency (congruent / incongruent) × amplitude (quartile bins 1–3). Other amplitude bin divisions (2, 4 bins) yielded similar results. The model was fitted using base R’s *aov()* with a fully crossed within-subjects error structure:

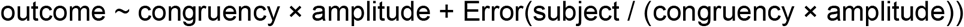

To control for multiple comparisons, FDR correction was applied separately for each experiment and dependent variable (183 tests per correction: 3 effects × 61 time windows).

For visualization purposes, we created time-amplitude heatmaps, where saccades were sorted into 10 amplitude bins. Bin edges were the deciles of the saccade-amplitude distribution computed from pre-sound saccades only. This kept the binning independent of any sound-evoked change in amplitude. Within each amplitude bin we run a running-window analysis with a 200 ms window and 50 ms step.

## Author Contributions

KJ: Conceptualization, Methodology, Software, Formal analysis, Data Curation, Investigation, Visualization, Writing - Original Draft; KB: Conceptualization, Methodology, Writing - Review & Editing; PB: Conceptualization, Software, Investigation, Writing - Review & Editing; ML: Conceptualization, Methodology, Writing - Original Draft, Supervision, Funding acquisition.

## Acknowledgement

The study was supported by the National Science Centre, Poland, OPUS 2022/45/B/HS6/04097.

## Supplementary Material

**Figure S1.**
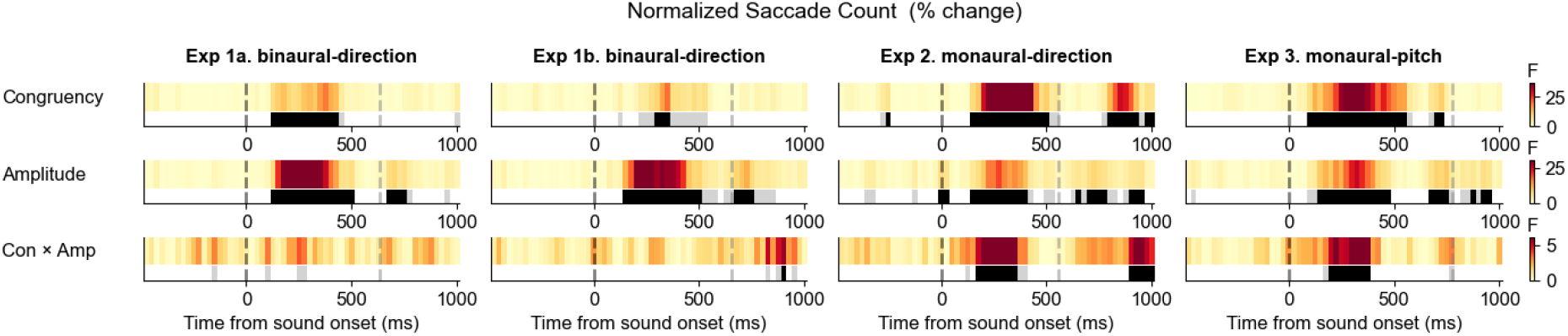
Effects of congruency and amplitude on normalized saccade count as a function of saccade timing. To examine the full time course of saccade amplitude and congruency effects on saccade frequency, repeated-measures ANOVAs were applied at each step of a 100 ms sliding time window, spanning −500 to +1000 ms, with 25 ms step (61 windows). Results are reported at both the uncorrected threshold (p < .05) in gray and the FDR-corrected threshold (p_FDR_ < .05) in black.

**Figure S2.**
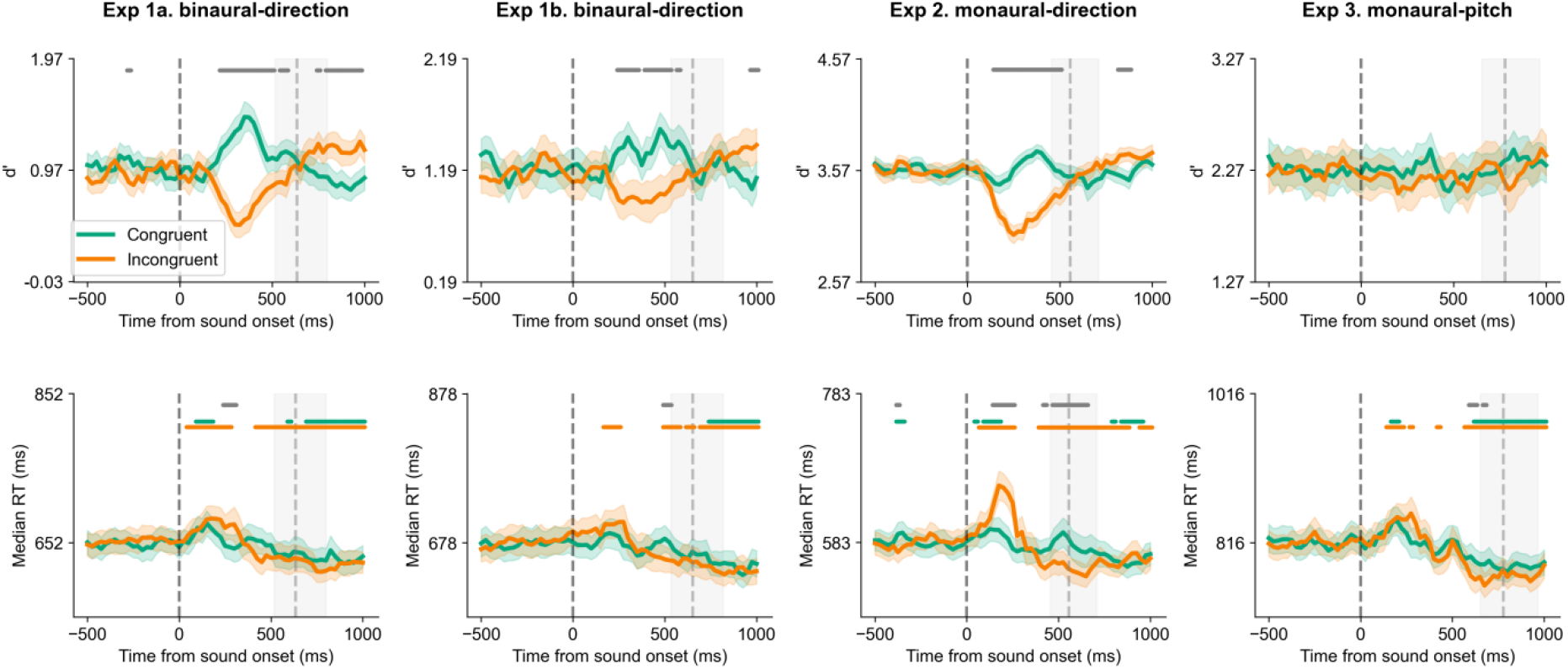
Other performance measures: d prime and reaction time, as a function of saccade timing. D prime and RT show the pattern of benefits and costs congruency with the probability of correct response measure. The Y axis was calibrated to show equal range around the mean value from the baseline, for all experiments. In all plots, shaded areas represent SEM across participants. Horizontal bars show significant, FDR-corrected difference from baseline for congruent (green) and incongruent (orange) saccades, and the difference between the relative count of congruent and incongruent saccades (gray). Note that for d prime statistical comparison relative to baseline was omitted (due to high volatility of the estimates with missing values and unequal samples, which occurred when comparing baseline to post-sound time windows). Black vertical dashed line indicates sound onset, gray vertical dashed line marks average manual response time in the auditory task, and gray vertical shading marks the 25%–75% range of response times.

**Figure S3.**
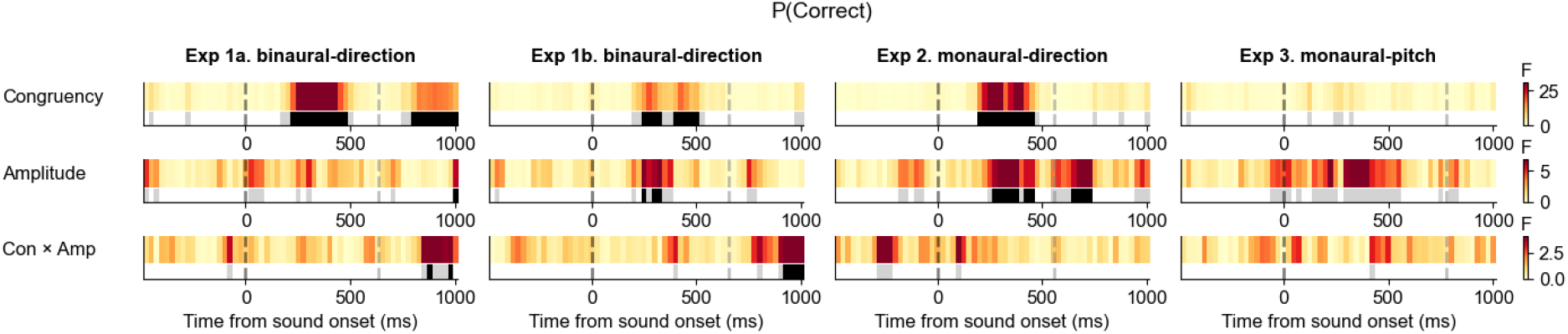
Effects of congruency and amplitude on the probability of correct response in auditory tasks as a function of saccade timing. To examine the full time course of saccade amplitude and congruency effects in auditory task performance, repeated-measures ANOVAs were applied at each step of a 100 ms sliding time window, spanning −500 to +1000 ms, with 25 ms step (61 windows). Results are reported at both the uncorrected threshold (p < .05) in gray and the FDR-corrected threshold (p_FDR_ < .05) in black.

